# Snekmer Learn/Apply: A kmer-based vector similarity approach to protein classification suitable for metagenomic datasets

**DOI:** 10.1101/2025.05.16.654600

**Authors:** Tara A. Nitka, Jeremy R. Jacobson, Christine H. Chang, Genevieve R. Krause, Travis J. Wheeler, Robert G. Egbert, William C. Nelson, Jason E. McDermott

## Abstract

Advances in whole genome sequencing have led to a rapid and ongoing increase in the amount of sequence data available, but 40-50% of known genes have no functional annotation and only 25-30% have specific functional annotations. Current functional annotation approaches typically rely on computationally expensive pairwise or multiple sequence alignments, preventing rapid development of models for novel protein functions and sometimes limiting methods to one ontology. Representation of sequence in short segments (kmers) has been used in many applications for nucleotide sequence, and more recently has been applied to protein sequence as well. We previously developed Snekmer, a tool which uses kmer patterns to develop alignment-free individual protein family models. Other approaches, such as MMSeqs2 and DIAMOND, use protein kmers as a fast filter to reduce search space for subsequent sequence alignment. Here, we describe a novel addition to the Snekmer tool which builds kmer libraries for protein families and uses those libraries to map functional annotations to new sequences. We first demonstrate that our method accurately applies TIGRFAMs annotations to protein fragments and to a low-sequence identity benchmark dataset, and further use it to annotate a set of drought stress associated soil and rhizosphere metagenome sequences with higher sensitivity towards several important protein function classes than that shown by HMMs. We have incorporated this workflow into Snekmer.

## Introduction

Advances in whole genome sequencing have led to the availability of a vast and rapidly increasing amount of sequence information. However, only 25-30% of microbial genes have specific functional assignments and 40-50% have no functional annotation at all (Lobb, et al., 2020; Salzberg, 2019). Current functional annotation methods typically rely on sequence similarity (e.g. BLAST (Altschul, et al., 1990; Altschul, et al., 1997), MMSeqs2 (Mirdita, et al., 2019; Steinegger and Söding, 2017), and DIAMOND (Buchfink, et al., 2021)) or hidden Markov models (HMMs). However, these methods are limited by both the inability of sequence similarity methods to reliably detect distantly related proteins and HMM methods are more computationally intensive for both building models and searching against them (Bateman, et al., 2000; Eddy, 1998). More recently, machine learning methods such as protein language models, a category that includes ProteinBERT (Brandes, et al., 2022), ProtTrans (Elnaggar, et al., 2022), and EAT (Elnaggar, et al., 2022) represent proteins as vectors of amino acids. While protein language models can annotate sequences not detected by sequence alignment methods, they struggle to distinguish decoy sequences from true hits to a family (Olson, et al., 2025) and are limited by high computational costs, particularly that associated with training. Some recent annotation tools such as MMSeqs2 (Mirdita, et al., 2019; Steinegger and Söding, 2017) and DIAMOND (Buchfink, et al., 2021) use short sequence motifs (kmers) to find the seeds they use to build alignments to improve speed, while RAST assigns annotations using “signature” kmers that occur in at least 80% of sequences within a family (Brettin, et al., 2015). Snekmer (Chang, et al., 2023), which was previously published by our group, combines the kmer approach with a reduced amino acid alphabet and vector representation of proteins.

Current functional annotation pipelines are limited by the computational cost of annotating large sequence datasets. Additionally, methods based on sequence similarity or HMMs are unable to effectively assign functional annotations to proteins which are detected and sequenced as fragments (Prakash and Taylor, 2012). Structure-based methods such as Foldseek (van Kempen, et al., 2024) depend on predicted structures, which can limit their effectiveness when applied to proteins with intrinsically disordered domains. Most existing kmer-based methods, such as RASTtk (Brettin, et al., 2015), eCAMI (Xu, et al., 2019), SimRank (DeSantis, et al., 2011), and kAAmer (Déraspe, et al., 2022) rely on exact kmer matches using the full amino acid alphabet, limiting k to small values due to the number of possible amino acids. Snekmer extends the kmer approach by incorporating multiple amino acid recoding (AAR) schemes to allow greater flexibility in sequence representation and capture relationships undetected by conventional methods (McDermott, et al., 2019). The resulting kmer vector representations of protein sequences are used to perform supervised model building and unsupervised clustering of protein families. This allows users to automatically build models for functional annotation and apply them to novel sequences, or to detect functional similarity by unsupervised clustering (Chang, et al., 2023). While both the AAR approach used by Snekmer and DIAMOND and MMSeqs2’s similar kmer approach have shown increased sensitivity relative to other kmer-based tools, we have previously shown that Snekmer and MMSeqs2 each capture a different set of otherwise-undetected relationships (Chang, et al., 2023). However, the original version of Snekmer is limited to smaller datasets than MMSeqs2, as memory requirements become excessive for large datasets, and it builds models for families individually which does not allow easy relative comparison of scores for query sequences. Finally, the original version of Snekmer builds models that are then fixed and have to be rebuilt to update, a feature shared by most family-based model methods.

Here, we describe a new Snekmer execution mode that allows rapid development of new model libraries spanning many thousands of protein families and application to hundreds of thousands to millions of sequences at a time with low computational cost (∼4x faster than HMMER). We present a new algorithm that allows rapid learning of protein family patterns at scale through association of annotations with kmer frequency, and probability-based scoring. The algorithm is incorporated into our existing Snekmer framework (version 1.3.0). We demonstrate that this method shows good (>90%) fidelity with HMM-based approaches but can be trained on new data much more easily and quickly, thus enabling rapid application to protein sequences. We also demonstrate that our method yields higher precision and recall than DIAMOND2, and is faster than DIAMOND2’s ultra-sensitive mode on small datasets. Furthermore, we demonstrate that our kmer-based approach can outperform HMMs for annotation of protein fragments, such as those often identified from short read sequencing of metagenomes. We demonstrate application of this method toward metagenomes from root rhizosphere communities, showing that our method can significantly improve the biological insight from such datasets with improved speed.

## Results

### Snekmer Family-wise Model Algorithm

The original Snekmer Model and Search workflow uses amino acid recoding and kmerization to train logistic regression classifiers for family assignment, it scores query sequences by individual family models, which makes comparison between hits digicult, and can produce biased results when some families are much longer than others. In addition, it is limited to small datasets by memory requirements, to address these limitations, we have developed a new Snekmer workflow, Learn and Apply, which uses the same Snekmer principles to simultaneously evaluate input protein sequences against a collection of protein families. In this new workflow, Snekmer builds models from training sequences by recoding, transforming these sequences into vectors of kmers, building a kmer count matrix which records the number of times each kmer occurs in each family, and then computes the cosine similarity of both the training sequences and ‘decoy sequences’ - reversed training sequences - to each family (see figure 1). The difference between cosine similarity of the training sequences to their correct families and their most-similar other family is used to generate a global confidence score by calibrating these scores with known family assignments, while the similarity of decoy sequences to each family is used to determine per-family cosine similarity thresholds, where a match to a decoy sequence is counted as a false positive.

**Figure 1.**
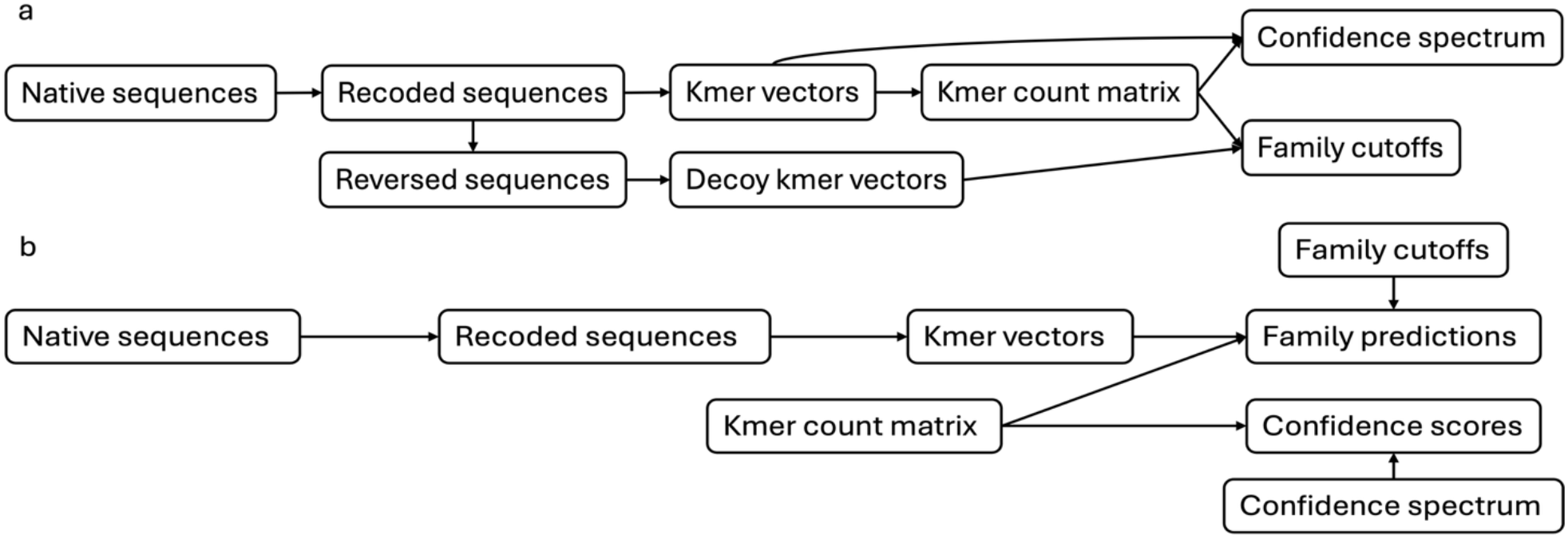
Generalized directed acyclic graphs of a) the Snekmer Learn workflow and b) the Snekmer Apply workflow.

### Snekmer multi-family evaluation on UniProt genomes

The efficacy of the Learn/Apply workflow was tested against a Hidden Markov Model approach. The TIGRFAMs (Tatusova, et al., 2016) set of protein families was used as the basis for this comparison, and HMMER (Eddy, 2011), which provides state-of-the-art sensitivity and specificity for protein family determination (Krause, et al., 2024; Steinegger and Söding, 2017) was the competing algorithm. For this assessment, 1,000 proteomes were randomly selected from UniProt, and an 80/20 train-test split (see table 1)was used to gauge the precision of various kmer sizes and amino acid recoding alphabets. Based on previous results (Chang, et al., 2023) three different amino acid alphabets were assessed: (1) a three-value alphabet based on solvent accessibility (SOLV), (2) a two-value alphabet distinguishing hydrophobic versus hydrophilic residues (HYD), and (3) a thirteen-value alphabet derived from the MIQS substitution matrix (Tomii and Yamada, 2016) (MIQS). Kmer length ranges tested were tailored to alphabet complexity: 7–9 for SOLV, 13–14 for HYD, and 3–4 for MIQS.

**Table 1.** TIGRFAM representation across training and test datasets derived from UniProt.

| Dataset | TIGRFAMs present | Proteins with TIGRFAM assignments (HMMER) | Proteins with TIGRFAM assignments (Snekmer) |
| --- | --- | --- | --- |
| Train | 3844 | 846,066 | 603,665 |
| Test | 3738 | 211,655 | 146,355 |

Notably, Snekmer Apply evaluation greatly outperformed HMMER in computational time. While HMMER (hmmsearch) required 6,960 core-minutes to search the 3784 TIGRFAMs HMMs against the 185,048 proteins in the test set, Snekmer completed the task in 480 core-minutes, demonstrating over a 14-fold increase in speed.

The best performing Snekmer Learn/Apply parameter combination in this test was SOLV and a kmer length of 8, which showed a precision of 90.7% and 90.7% recall, using HMMER family assignments as the gold standard for this analysis. Screening Snekmer results for those with a confidence score >0.95 eliminated predictions for 35,105 sequences (19% of total) and increased precision to 98.7% while decreasing recall to 79.9%.

To determine whether Snekmer’s performance is affected by structural or functional properties of a family, results from the above analysis were re-evaluated, grouping families by protein length, sequence complexity (low entropy, high entropy), localization (cytosol, membrane, extracellular), and function (enzyme, non-enzyme, highly conserved functions such as DNA repair, translation or ribosomal proteins, or variable functions like transporters) (see Methods). We also examined results with families grouped by the number of training sequences. In this analysis, Snekmer showed >90% accuracy across most categories (Figure 2), but lower accuracy was observed for low entropy sequences, transmembrane proteins, and transport proteins. Accuracy is also sensitive to the number of sequences in the training data. The lower accuracy in these cases is likely the result of the high number of repeats in low entropy sequences, which is a known challenge for functional annotation (Tørresen, et al., 2019), and the possibility that families with small numbers of representative sequences result in non-representative family cutoff scores.

**Figure 2.**
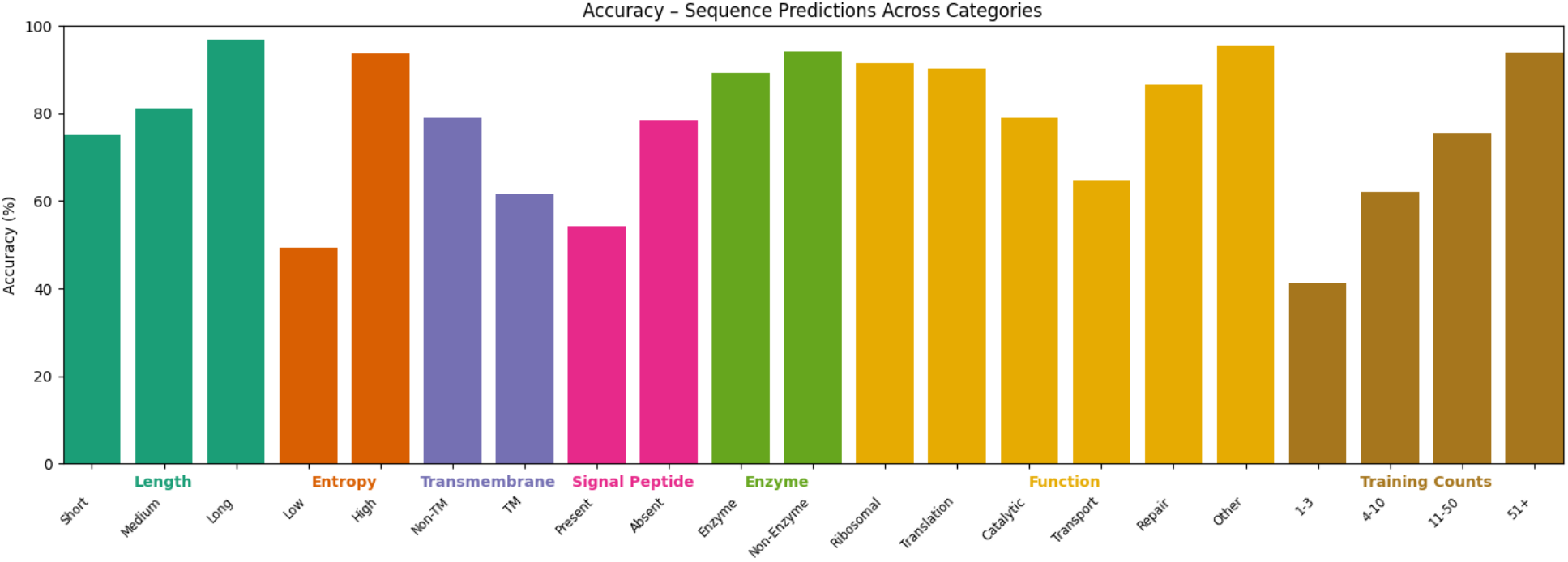
Comparison of Snekmer accuracy by sequence length, sequence entropy, presence of transmembrane domains, presence of signal peptides, function, and number of sequences in the correct family’s training data.

### Snekmer shows increased sensitivity for fragmentary data

We hypothesized that Snekmer, as a kmer-based approach, would have greater sensitivity against fragmentary input data than methods optimized for full-length sequences, such as TIGRFAM HMMs and alignment-based tools. This was tested by generating a test data set of artificial fragments with varying length (25-200 amino acids) by randomly sampling sections of the full-length sequences from 200 UniProt proteomes. These fragments with TIGRFAMs families using both Snekmer and HMMER, and compared them against the annotations determined by application of HMMER to the full length proteins. While HMMER retains the high precision shown on full length sequences, Snekmer has superior recall on all fragments tested and has precision >0.8 on fragments of 125 residues or longer. The precision and recall of Snekmer and HMMer are displayed in Figure 3.

**Figure 3.**
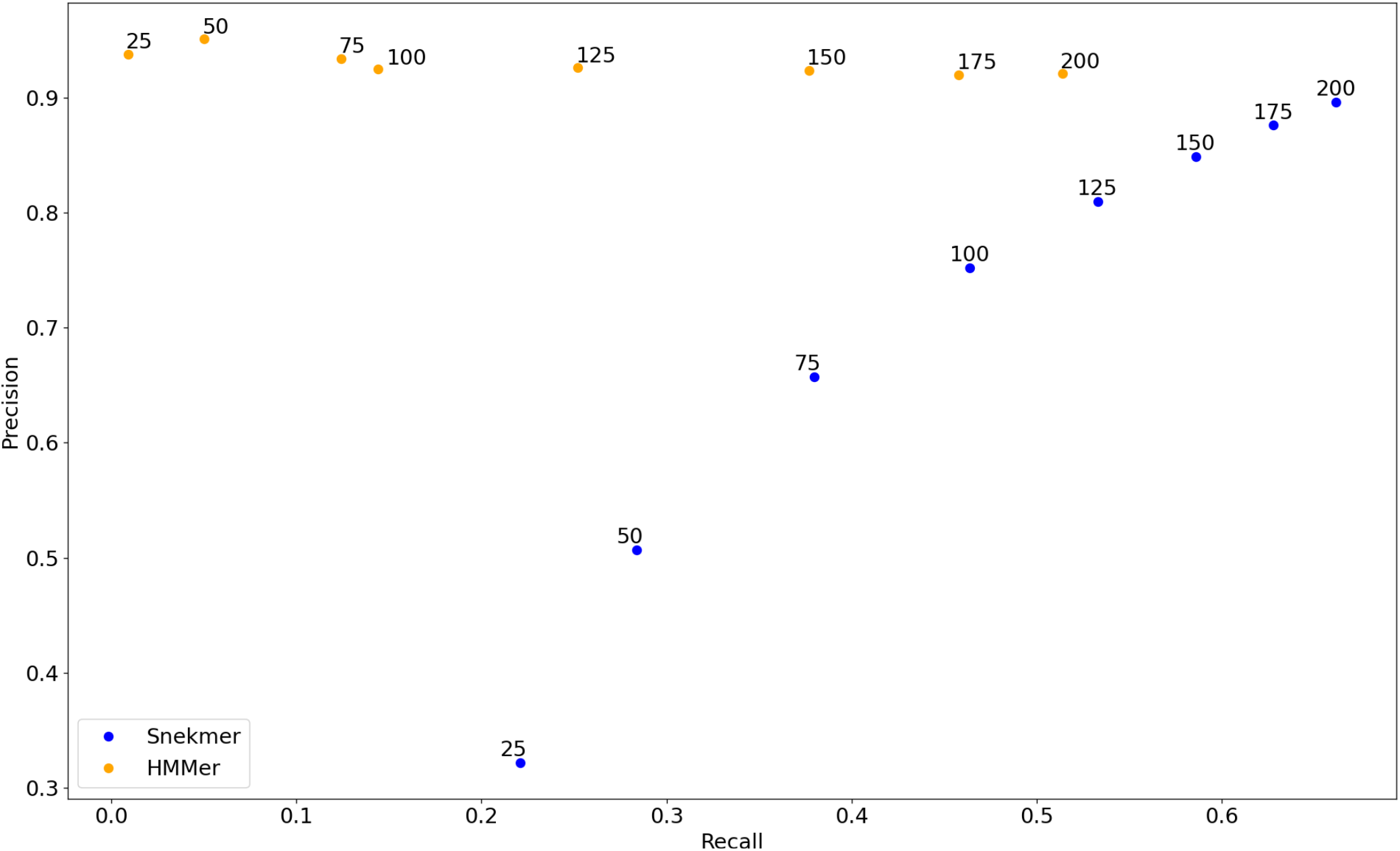
Precision vs. recall of TIGRFAMs functional annotations applied to sequence fragments by Snekmer and HMMER. Points are labeled by sequence length.

### Snekmer phylogenetic sensitivity

To determine Snekmer’s sensitivity across evolutionarily distant sequences, we examined Snekmer’s performance when trained on a set of genomes that were not in the same evolutionary family as the target sequences. The set of 1000 UniProt genomes was divided into seven well-represented clades (Alphaproteobacteria, Bacilli, Bacteroidota/Chlorobiota, Betaproteobacteria, Clostridia, Gammaproteobacteria, or Pseudomonadota) totaling 891 genomes. Then a leave-one-out analysis was performed where the set of genomes from each clade in turn was excluded from the training data, and performance of the resulting models was evaluated (Table 2).

**Table 2.** Results of Snekmer phylogenetic leave-one out analysis. Results are filtered using a 95% confidence threshold.

|  | Test genomes (number of proteins) | Precision | Recall |
| --- | --- | --- | --- |
| Bacteroidetes | 111 (95,684) | 97.36% | 54.72% |
| Pseudomonadota | 353 (350,082) | 96.88% | 58.39% |
| Betaproteobacteria | 71 (77,257) | 98.85% | 70.32% |
| Gammaproteobacteria | 132 (127,955) | 98.64% | 67.57% |
| Alphaproteobacteria | 143 (141,748) | 98.42% | 63.27% |
| Clostridia | 78 (69,729) | 97.78% | 68.78% |
| Bacilli | 101 (95,800) | 97.89% | 68.89% |

As another assessment of sensitivity across evolutionary distance, we performed a similar analysis where a model set was trained on sequences from only one clade and applied to the other bacterial clades. For example, using a model trained only on the Betaproteobacteria, we observed lower precision, ranging from 66.6% to 93.3% across the other clades (Figure 4). Recall ranged from 65.4% to 92.4%, increasing inversely with evolutionary distance (see supplementary figure S4) as measured by estimated divergence time between clades retrieved from TimeTree (Kumar, et al., 2017). This pattern indicates that sensitivity is reduced for distantly related sequences but precision remains high after confidence filtering.

**Figure 4.**
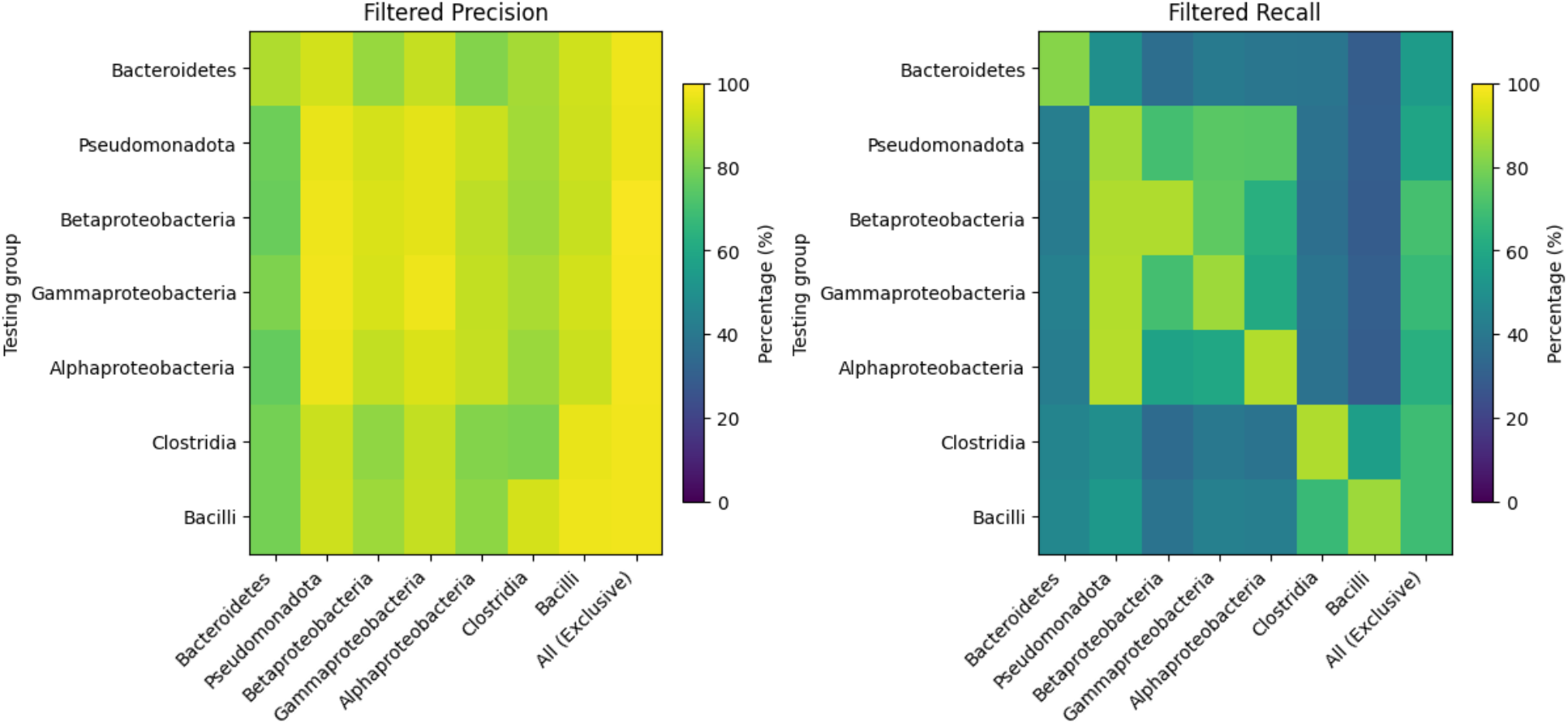
Precision and recall of Snekmer annotations using a training set containing genomes from only one clade. Results are filtered using a 95% confidence threshold. The rightmost column of each heatmap shows the results when applying a model to all clades except that used for training.

### Benchmarking Snekmer in the sequence similarity ‘twilight zone’

The above results demonstrate that Snekmer can reasonably recapitulate HMMER results for the ‘easy’ case where evaluation sequences are potentially similar to those used for training, and that Snekmer can perform well across moderate evolutionary distance between organisms. To assess Snekmer’s ability to classify sequences that are significantly different than those in the training set (related by less than 25% sequence identity, the so-called ‘twilight zone’ (Doolittle, 1986; Rost, 1999)), we created rigorous benchmark training and test data sets using the profmark approach (Eddy, 2011) to split the seed alignments for TIGRFAMs HMMs into training and test sets. Sequences were split using the single-linkage clustering algorithm integrated into profmark, such that no test sequence has more than 25% identity to any training sequence or more than 50% identity to another test sequence. Training and test data sets were successfully created for 341 TIGRFAMs. We evaluated Snekmer and HMMER results on this dataset using receiver operating curves where false positive rate is the fraction of decoys annotated and true positive rate is the fraction of correct test sequence annotations. Test sequences that were classified into multiple families were omitted from this analysis because they may truly be homologous to more than one family. Using this approach to evaluate Snekmer using the 3 previously used AAR alphabets as well as the native 20 amino acid alphabet (native), a 7-value chemical property alphabet (standard), and a 3-value alphabet based on HYD with charged residues as a third category (hydrocharge), with a variety of kmer lengths, we found that the solvent accessibility alphabet and k=8 was the best compromise between sensitivity and specificity (AUC-ROC = 0.89); other combinations of alphabet and k yielded AUC-ROCs ranging from 0.59 to 0.85 (Figure 5). In comparison, the AUC-ROC for HMMER using the HMMs built from the profmark training alignments was 0.99.

**Figure 5.**
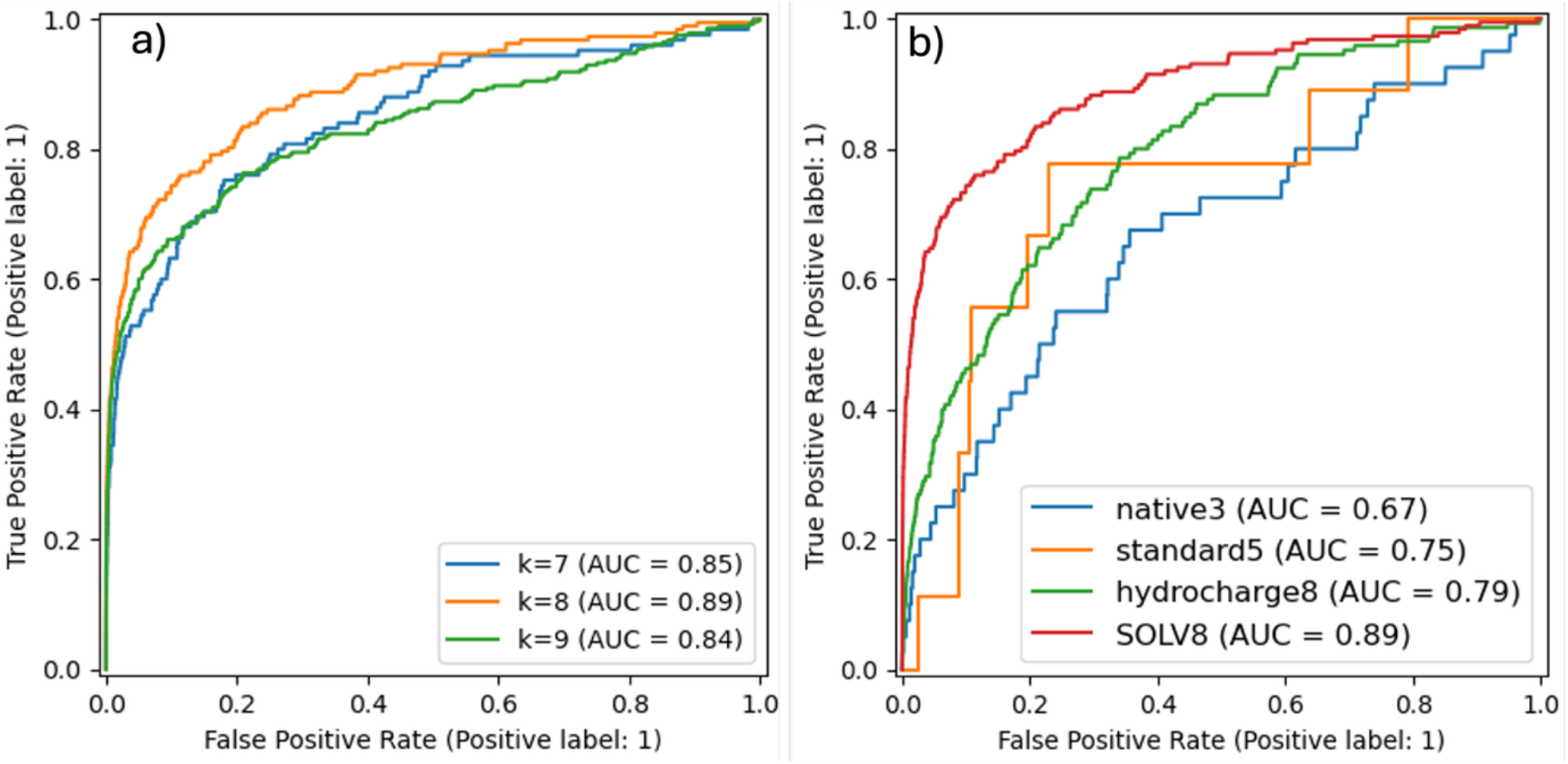
Performance results for profmark testing. a) ROC plot showing Snekmer performance using the solvent accessibility alphabet and k between 7 and 9; b) ROC plot showing Snekmer performance for the best-performing k associated with 5 amino acid recoding alphabets and the native 20 amino acid alphabet, the 3-value hydrophobicity/structure breaker alphabet is omitted because it had much lower coverage than any other alphabet.

These results show that, using the best combination tested, Snekmer underperforms HMMER in both sensitivity and specificity. However, Snekmer’s performance in this very digicult evaluation is reasonable, and indicates that it will be useful for making predictions for this kind of ‘dark matter’, with the caveat of reduced confidence for these predictions.

### Application of Snekmer to soil MAGs

To demonstrate Snekmer’s performance on a real-world dataset, we annotated a set of 244,087 predicted proteins from 55 previously published metagenome-assembled genomes (MAGs) from soil and sorghum rhizosphere samples (Xu, et al., 2021). Compared to DIAMOND and HMMER, Snekmer showed higher annotation coverage, annotating 66.5% of sequences at a minimum of 95% confidence compared to the 16.8% annotated by HMMER. Snekmer was also substantially faster when compared to HMMER on a subset (43) of these MAGs (∼4.02x), requiring only 29.6 core-hours to annotate this dataset compared to the 119 core-hours needed using hmmsearch. DIAMOND2 required 39.7 core-hours for these MAGs, but yielded only 60.0% precision and 24.2% coverage.

The higher coverage of Snekmer annotations relative to TIGRFAMs HMMs is primarily driven by higher coverage of proteins involved in the general KEGG metabolic categories of biosynthesis, ‘Energy metabolism’, ‘Transcription’, ‘Cell motility and adherence’, and ‘Defense and invasion systems’ (Figure 6). Snekmer also made more assignments to TIGRFAM families whose exact function is unknown.

**Figure 6.**
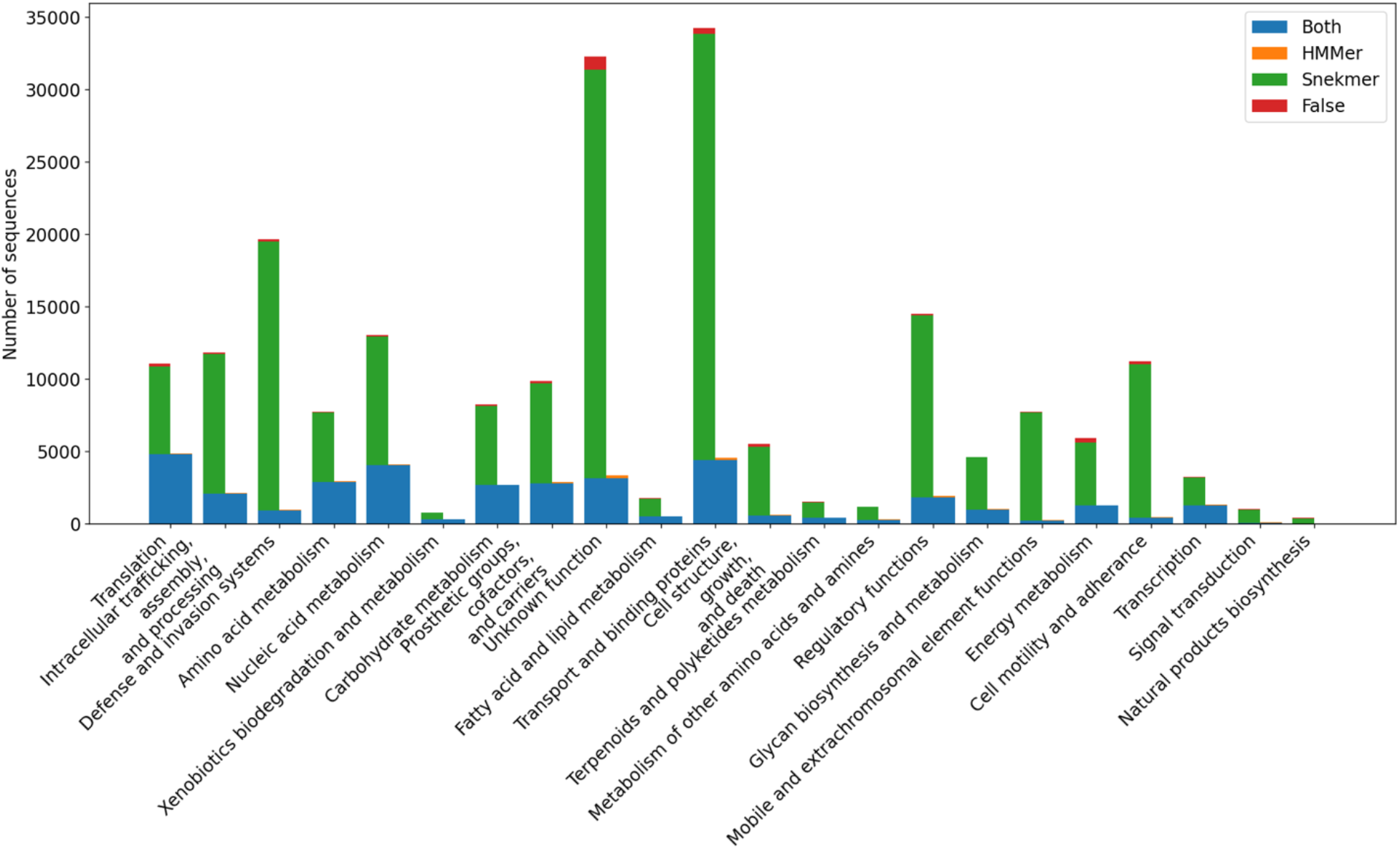
Functional role annotations by HMMER and Snekmer. Annotations assigned by both methods are shown in blue, the expected number of false positives is shown as a red section of the bar showing annotations unique to Snekmer (HMMER is expected to have 0 false positive annotations).

To verify that Snekmer’s increased annotations relative to HMMER do not stem from false positives, we repeated the analysis using reversed sequences (Glidden-Handgis and Wheeler, 2024) from the drought MAGs. On this reversed dataset of 244,087 sequences, Snekmer produced 3,151 false positive annotations (FPR = 1.3%), while HMMER produced none. Given that the number of false positives on the reversed sequences is small relative to the 162,409 annotations assigned to the forward sequences, we attribute Snekmer’s higher coverage on the drought MAGs to be primarily from increase in sensitivity toward certain protein families rather than a tendency for false assignments (Figure 6). We performed the same analysis using DIAMOND2, transferring annotations from the best-scoring alignment target reported using the default e-value cutoff of 10^-3^, which produced 129 false positive annotations (FPR = 0.1%) but yields coverage between that of HMMER and Snekmer on the forward sequences (Supplementary Figure S1).

### Conclusions and future directions

We describe our new Snekmer workflow, Learn/Apply, which we have developed to provide faster model development and searches than the existing Model/Search workflow. Comparison of this method to HMMER and DIAMOND2 shows that it provides comparable precision to HMMER with higher metagenomic coverage and lower computational cost, while providing greater coverage and precision than DIAMOND2. Because of the self-evaluation method used to determine the quality of Learn/Apply annotations, the family cutoff scores are less stringent than those typically used in HMMER. This can be beneficial, as it allows higher annotation coverage of protein fragments, and allows sequences with functions that lack a specific model to be assigned to the most-similar available family. However, as the latter behavior is sometimes undesirable, Model/Search or HMM-based techniques may be needed for applications where specificity is critical. However, Learn/Apply can still reduce the time needed to identify proteins with a highly specific function, as metagenomic data can be rapidly annotated with Learn/Apply to allow using Model/Search or HMMs to search only the sequences assigned to the family of interest rather than searching an entire metagenome with these techniques. Because Learn/Apply outperforms HMMER for annotation of protein fragments, it can be used either as a standalone tool or alongside another fast method such as MMSeqs2 to rapidly annotate sequence fragments with higher coverage and lower computational cost than HMMER.

By representing proteins and protein families as kmer count vectors and comparing those vectors using cosine similarity, the new Snekmer Learn/Apply workflow increases the scale and speed of functional assignment to protein sequences with lower computational complexity and higher coverage than TIGRFAMs HMMs. Further, we have demonstrated that the AAR/kmer vector approach implemented in Snekmer shows only a small decrease in performance relative to HMMER on sequences from evolutionarily distant bacteria while requiring lower computational cost and it achieves higher coverage for annotation of fragmentary sequence data. Snekmer’s higher sensitivity on fragmentary data is of particular importance, as high-throughput sequencing techniques produce short reads, and sequences assembled from short read sequencing data, especially from metagenomic samples, typically generate a significant number of protein fragments.

While the per-residue information content is lower for a recoded sequence, the results on the profmark benchmark show that Snekmer’s AAR approach accounts for sequence diversity beyond that present in the training data. Because AAR captures greater flexibility in sequence space (McDermott, et al., 2019; Tomii and Yamada, 2016), family models are able to recognize sequences as members of a family even when they contain a number of conservative substitutions that could cause them to be missed if AAR were not used. The phylogenetic drop-out analysis shows that this also allows orthology to be detected across different clades. However, there is still an inverse relationship between evolutionary distance and precision/recall. Because this relationship likely results from substitutions in positions with low conservation, which are likely less important to protein function, it is possible that careful selection of the AAR alphabet used could increase cross-taxa precision and recall for a family of interest. Development of AAR alphabets based on sequence conservation may also improve cross-taxa performance, although the current MIQS alphabet does not perform well on the profmark benchmark. However, this may be because of the smaller kmers necessitated by larger alphabets, which suggests the possibility of improvement through further optimization of Snekmer to reduce the compute and memory requirements associated with large AAR alphabets.

Because Snekmer annotations, unlike HMMER, do not directly depend on sequence length, Snekmer shows much higher recall than HMMER when tested on sequence fragments. Snekmer’s accuracy is still reduced for short fragments, but is at least 0.80 for fragments of 125 residues or longer and approaches HMMER’s accuracy on fragments of 200 residues, while Snekmer’s advantage in recall also improves with fragment length. The better performance shown by Snekmer likely results from a combination of the kmer approach and the use of cosine similarity, which allows sequences to be assigned to a family if enough representative kmers are present without depending on the total number of kmers present.

The dramatic increase in coverage demonstrated for the soil MAGs (Figure 6) over what is annotated by HMMER is surprising. We looked into these extra annotations and found a number of features indicating that they are reasonable predictions made by Snekmer.

Though the number of decoy sequences matched by Snekmer was not zero, it was a relatively small percentage indicating that only slightly over 1% of the novel annotations (nearly 60,000) is likely due to inappropriate matches. Examining the novel predictions for the defense and invasion systems functional group, we found that many of them were ATP-binding proteins within ABC transporter complexes and type III secretion system proteins with functions closely related to the TIGRFAM predicted by Snekmer, but not actually members of any TIGRFAM. This confirms that Snekmer is, as we expect, assigning sequences to TIGRfams which are closely related to their function. While these sequences are not truly members of any TIGRfam, these annotations indicate that Snekmer reliably finds sequence orthology and provides evidence of function even for sequences whose exact function is not represented by any model used. This suggests that most apparent “false positive” annotations are the result of orthology between a protein family not represented by the models being used and a model for a similar family, rather than uninformative matches. Finally, it is worth noting that the HMMER TIGRFAMs model searches we performed used trusted bit score cutoffs for each family that are based on the full length of the protein, which makes the matches more conservative and likely less sensitive to fragmentary data. Interestingly, the DIAMOND2 searches, which do not include a conservative trusted cutoff and instead use an e-value cutoff of 10^-3^, yield very similar results to the HMM results. Though further investigation is needed, we have shown that the additional annotations provided by Snekmer in a real-world application are reasonable and could greatly enhance the annotation of partial genomes and metagenomes.

Snekmer shows high accuracy and recall across protein classes, performing best on enzymes and high sequence entropy proteins. While Snekmer does show lower accuracy on low sequence entropy proteins, transmembrane proteins, and families with few training sequences, this is likely to be the result of repetitiveness in low sequence entropy and transmembrane proteins, which is an issue for many annotation methods (Tørresen, et al., 2019). The problem of small training data likely indicates that the value of family-specific cutoff scores is dependent on the size of training data, and can be avoided by including at least 30 sequences in each family if adequate data is available. Low numbers of available target sequences are also likely to pose an issue for other methods, as they are unlikely to represent the full sequence diversity of a protein family.

The Snekmer TIGRFAM models show much higher metagenomic coverage than the corresponding HMMs, particularly for proteins involved in biosynthesis, energy metabolism, transcription, cell motility and adherence, and defense and invasion. Snekmer also shows higher sensitivity towards iron uptake proteins, which are relevant to the iron metabolism perturbation observed by Xu *et al*. under drought stress conditions. Specifically, *Streptomyces* is among the most highly enriched genera under drought stress, and exhibits a DNA transfer process that is dependent on ATP hydrolysis, which Snekmer shows high coverage towards (Fitzpatrick, et al., 2018; Xu, et al., 2018).

Snekmer is also significantly less computationally intensive than HMMER, by a factor of 4.02 (29.6 vs 119 core-hours) on a 43-member subset of the 55 drought stress MAGs. Although the computational cost on large datasets is still higher than that of DIAMOND2, Snekmer shows higher coverage than DIAMOND2 with precision close to that of HMMs. It’s likely that Snekmer was faster on this dataset due to a combination of the facts that DIAMOND2 is optimized for query datasets of >1 million sequences, that DIAMOND2 was run in ultra-sensitive mode, and that the same sequences used to train Snekmer were used as a database for DIAMOND2, necessitating many more similarity comparisons by DIAMOND2 than by Snekmer. We expect DIAMOND2 to run faster than Snekmer on larger datasets or with a more carefully curated target database, but the difference in coverage and precision would not be affected by these factors. We have also previously shown that Snekmer’s amino acid recoding approach captures a different region of sequence space than MMSeqs2’s similar kmer approach (Chang, et al., 2023). Because Snekmer has nearly the same precision as HMMs with better coverage and computational performance, adding Snekmer to an annotation pipeline is very likely to improve results without compromising speed, even if that pipeline already contains methods such as MMSeqs2.

The Snekmer Learn/Apply workflow is a valuable complement to high-throughput annotation pipelines, as it can annotate sequences missed by other methods with high precision and low computational cost and allows the rapid creation of models representing novel functions. The Snekmer approach shows near-HMM precision with higher annotation coverage, particularly for sequence fragments, and lower computational cost. The use of Snekmer in combination with MMSeqs2 is particularly valuable because both the AAR approach used by Snekmer and the similar kmer approach used by MMSeqs2 detect relationships not found by each other or by HMMs, while also allowing much faster annotation than HMMs with only a small decrease in precision. Combining Snekmer Learn/Apply with a more selective method such as HMMs and the older Snekmer Cluster workflow could also provide a way to rapidly identify proteins that are similar to, but not members of, a known protein family, cluster those sequences into one or more novel families, and build a model to detect other members of those families.

## Methods

### Snekmer multi-family workflow

Snekmer’s multi-family function modes, called Learn and Apply, are built off the same theory and concepts as Snekmer’s other features, all of which utilize kmerization coupled with amino-acid recoding to retain positional information and represent the properties of amino acids while reducing the space complexity. As implied by their names, Snekmer multi-family modes represent paired functions which allow users to build a library that ‘learns’ how kmers are associated with protein families and then ‘applies’ them to new proteins.

The multi-family Learn function is a multi-step process embedded in a Snakemake pipeline. Initially, protein sequences are recoded using a user-defined amino acid recoding (AAR) scheme. These recoded sequences are then transformed into vectors of overlapping kmers (“kmerized”), where the length of each kmer (k) is a user-definable value. Next, the occurrences of each unique kmer are counted for each protein family to create a kmer count vector for each family. The count vectors for each family are then aggregated into a kmer count matrix. Two inputs are required for Learn: FASTA files for training, and an annotation file with sequence IDs mapped to protein function IDs.

Because families consisting of long proteins are likely to have less sparse kmer vectors, which may lead to cosine similarity scores being affected by non-informative kmers, family-specific thresholds are assigned using reversed input sequences (‘decoys’). After building the kmer count matrix, the family-specific thresholds are assigned by reversing, recoding, kmerizing, and vectorizing the input sequences, then computing the cosine similarity of the resulting sequence vectors to each family vector, and the median cosine similarity of these sequences to a family is used as the cutoff for assignment to that family.

Finally, to provide a global measure of classification precision, a confidence spectrum is calculated by self-evaluation. During self-evaluation, each training sequence is kmerized, vectorized, and compared to each family vector using cosine similarity. The difference in score between the true annotation and highest-scoring false annotation is calculated using equation 1. The resulting distribution is then used to calculate confidence scores using equation 2. Because the self-evaluation process is dependent on the number of training sequences, the use of small training datasets should be avoided as it can lead to overfitting and inaccurate confidence scoring.

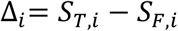

Equation 1. For each protein *i*, Δ_I_ is the difference between the top two cosine similarity scores, S_T,I_ is the highest similarity score (which is always the correct family during training) while S_F,I_ is the second-highest score.

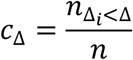

Equation 2. The confidence score for each value of Δ (c_Δ_) is the number of sequences with Δ_i_<Δ (n_Δi<Δ_) divided by the total number of input sequences (*n*).

Although Snekmer is optimized for processing all input data in a single instance, it can accommodate additive data integration. This approach yields an identical kmer count matrix, but the resultant confidence spectra may exhibit marginally reduced precision (Supplementary Figure S1).

The Snekmer kmer count associations, confidence spectrum, and family-specific thresholds used in this paper were generated using the Learn algorithm on 800 genomes obtained from UniProt (The UniProt Consortium, 2022).

The multi-family Apply function operates on three inputs: the kmer count matrix, its corresponding confidence look-up table produced during Learn, and a collection of protein FASTA files. First, the Apply workflow recodes, kmerizes, and vectorizes input sequences utilizing the same methodology as Learn, then calculates the cosine similarity between each vectorized sequence and each family vector stored within the kmer count matrix.

Cosine similarity is used because it depends only on the direction of the sequence vector, not its magnitude, which facilitates better performance toward sequence fragments as sequence length does not affect the resulting annotation. Cosine similarity scores are then compared to the family-specific thresholds, and hits which do not meet the threshold are discarded. The remaining family with the highest cosine similarity score becomes the annotation for that sequence.

The code for both Learn and Apply is available as part of Snekmer, at https://github.com/PNNL-CompBio/Snekmer.

### Comparison to HMMER and DIAMOND2

HMMER analyses were performed using TIGRFAMs version 15 (Li, et al., 2021) with HMMER version 3.3.2 (Eddy, 2011). The TIGRFAMs trusted cutoffs were used for this analysis.

DIAMOND2 analyses were performed using DIAMOND version 2.1.8.162 (Buchfink, et al., 2021). The same 800 genomes used to train Snekmer were used to build a DIAMOND2 database, and searches were performed in ultra-sensitive mode. Annotations were assigned by taking the best-scoring alignment for each sequence and transferring the target protein’s TIGRFAM annotation to the query sequence.

Apply and DIAMOND2 results were compared to HMMER results using Python version 3.9.14. The BioPython library (Cock, et al., 2009) was used to parse HMMER tabular output files, data was assembled into numpy arrays (Harris, et al., 2020), and other data handling was done using Pandas (Mckinney, 2010; The pandas development team, 2024).

### Performance evaluation

Performance of each tool was evaluated by calculating the precision and coverage or recall, as appropriate, on each dataset. Coverage is defined as the percentage of sequences receiving annotations. For precision and recall calculations, we define true positives (TP) as annotations matching that assigned by HMMER, and false negatives (FN) as any sequence which is missed by Snekmer despite receiving a HMMER annotation. False positives (FP) are defined as annotations assigned to decoy sequences. Any sequence receiving annotations from neither Snekmer nor HMMER is considered a true negative (TN). Snekmer annotations to positive test sequences that differ from the annotation assigned by HMMER are excluded from calculations, as it is uncertain whether the sequence has some relationship to that family or is simply incorrect. Precision is calculated using the equation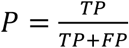, while accuracy is calculated as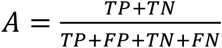 and recall is calculated using the equation 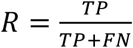.

### Data sources

The TIGRFAMs version 15.0 HMMs were retrieved from NCBI using FTP (https://ftp.ncbi.nlm.nih.gov/hmm/TIGRFAMs/release_15.0/) and compressed and indexed using hmmpress. A set of 1000 randomly selected bacterial genomes was obtained from UniProt (The UniProt Consortium, 2022).

Sequence data for the drought stress soil and rhizosphere MAGs was provided by Xu et al. (Xu, et al., 2021). Gene calling and translation of the provided data were performed using Prodigal (Hyatt, et al., 2010), running in “normal” (single-genome, training using input data) mode for drought stress MAGs.

Estimated divergence times between classes used in the phylogenetic sensitivity analysis were obtained using TimeTree (Kumar, et al., 2017).

### Snekmer benchmarking v. HMMER for the sequence similarity ‘twilight zone’

The benchmark dataset for comparison of Snekmer to HMMER 3.3.2 was constructed from the TIGRFAMs version 15 seed alignments using the profmark benchmark creation tool (Eddy, 2011). TIGRFAMs alignments were split into query alignments and test domains by two single-linkage clustering steps, with all test sequences required to have <25% sequence identity to any query sequence and downsampled such that no two test sequences share more than 50% sequence identity. Positive test sequences were constructed by profmark by embedding two test domains in a larger sequence with nonhomologous segments generated by shuffling a random protein segment from SwissProt. Negative test sequences were constructed by randomly sampling the segment length structure of positive test sequences and replacing all segments with nonhomologous shuffled segments. All benchmark runs used profmark’s master Perl script for batch submission of SLURM jobs, while the hmmsearch driver script and our own Snekmer driver script were used for HMMER and Snekmer benchmarks, respectively. The hmmsearch benchmark script uses the query alignments to construct profile HMMs, and then uses hmmsearch to search the test set for hits to each HMM. The Snekmer benchmark script uses the Learn mode to build kmer count associations and confidence scores for each query (family) alignment, then uses the Apply mode to calculate cosine similarity of each test sequence to each family. The profmark master script was modified only as needed to format SLURM scripts for our HPC cluster, while the hmmsearch benchmark driver script was used without modification. Both hmmsearch and Snekmer benchmarks were performed on a single 64-core node with 2.5 GHz clock rate and 256 GB available RAM. Code used to run this analysis is available at [TBD].

### Amino Acid Recoding

Snekmer supports amino acid recoding using 5 alphabets: a 2-value alphabet classifying residues as either hydrophobic or hydrophilic (HYD); a 3-value solvent accessibility-based alphabet (SOLV); a 7-value chemical property-based alphabet (standard); a 13-value alphabet based on the MIQS substitution matrix (MIQS); and a pair of 3-value alphabets that extend the hydrophobicity alphabet by adding a category for either charged residues (hydrocharge) or structure-breaking residues (hydrostruct). The mapping of each canonical amino acid to each recoding alphabet is shown in Table 3.

**Table 3.**
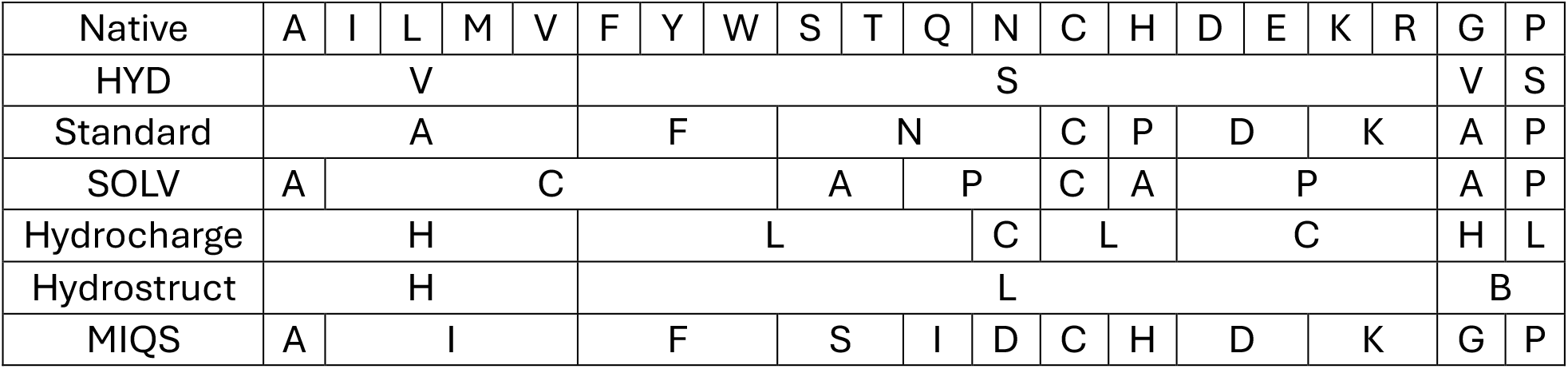
Amino acid recoding alphabets mapped to the native 20 amino acids. Note that “standard” refers to the standard 7-value chemical property alphabet, while the canonical amino acid alphabet is referred to as “native”.

## Funding

This work was supported by the US Department of Energy Biological and Environmental Research Division (DOE/BER) under the “Persistence Control of Engineered Function in Complex Soil Microbiomes” Science Focus Area (SFA) and the US Department of Agriculture Animal and Plant Health Inspection Service (USDA APHIS) project “A systems approach to understanding farm animal-environmental drivers of SARS-CoV-2 transmission in the food supply chain”. Pacific Northwest National Laboratory is operated by Battelle Memorial Institute for the US Department of Energy under contract DE-AC05-76RL01830.

## Conflict of Interest

None declared.

---

**Supplementary Figure S1.**
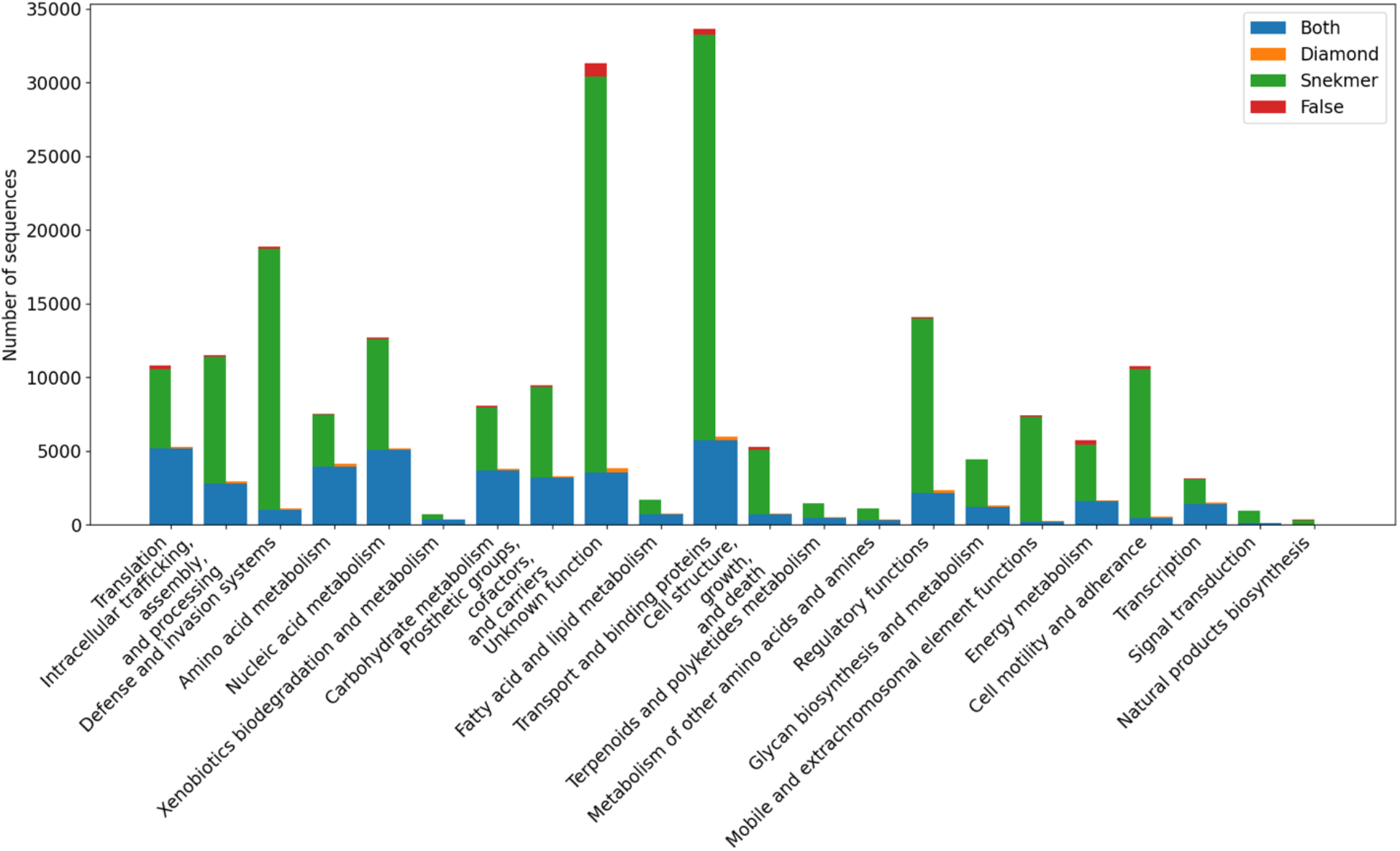
Functional role annotations by Snekmer and DIAMOND2. Annotations assigned by both methods are shown in blue, the expected number of false positives is shown as a red section of the bar showing annotations unique to Snekmer.

**Supplementary Figure S2.**
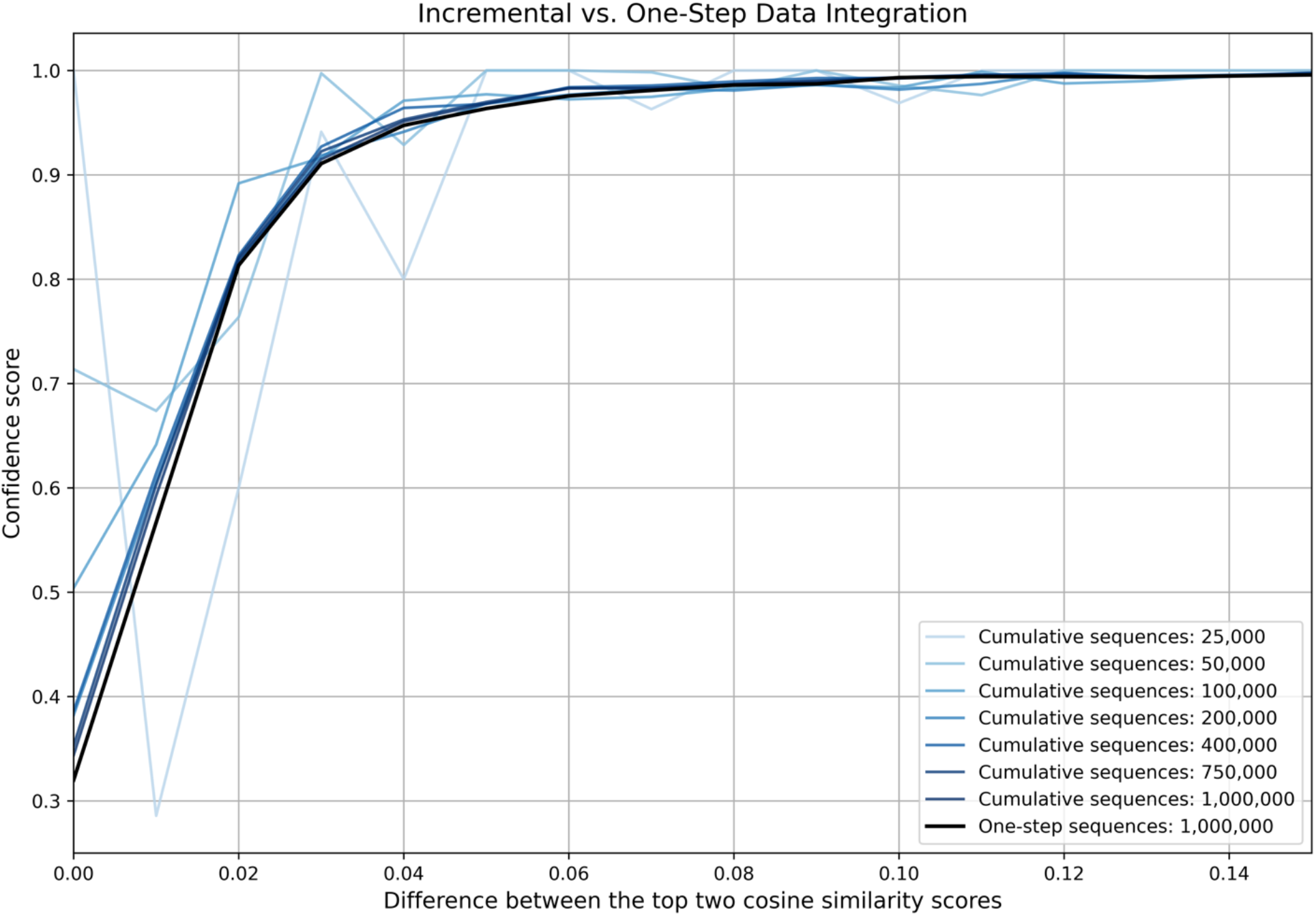
Confidence scoring using the additive approach vs single integration for 1 million sequences across 275 genomes.

**Supplementary Table S3.** Precision of Snekmer annotations on control sequences.

| Control Precision | Control Precision (filter: 95%) |
| --- | --- |
| 0.9080 | 0.9902 |
| 0.9038 | 0.9896 |
| 0.9041 | 0.9882 |
| 0.9088 | 0.9903 |
| 0.9044 | 0.9887 |
| 0.9014 | 0.9898 |
| 0.8927 | 0.9888 |

**Supplementary Figure S4.**
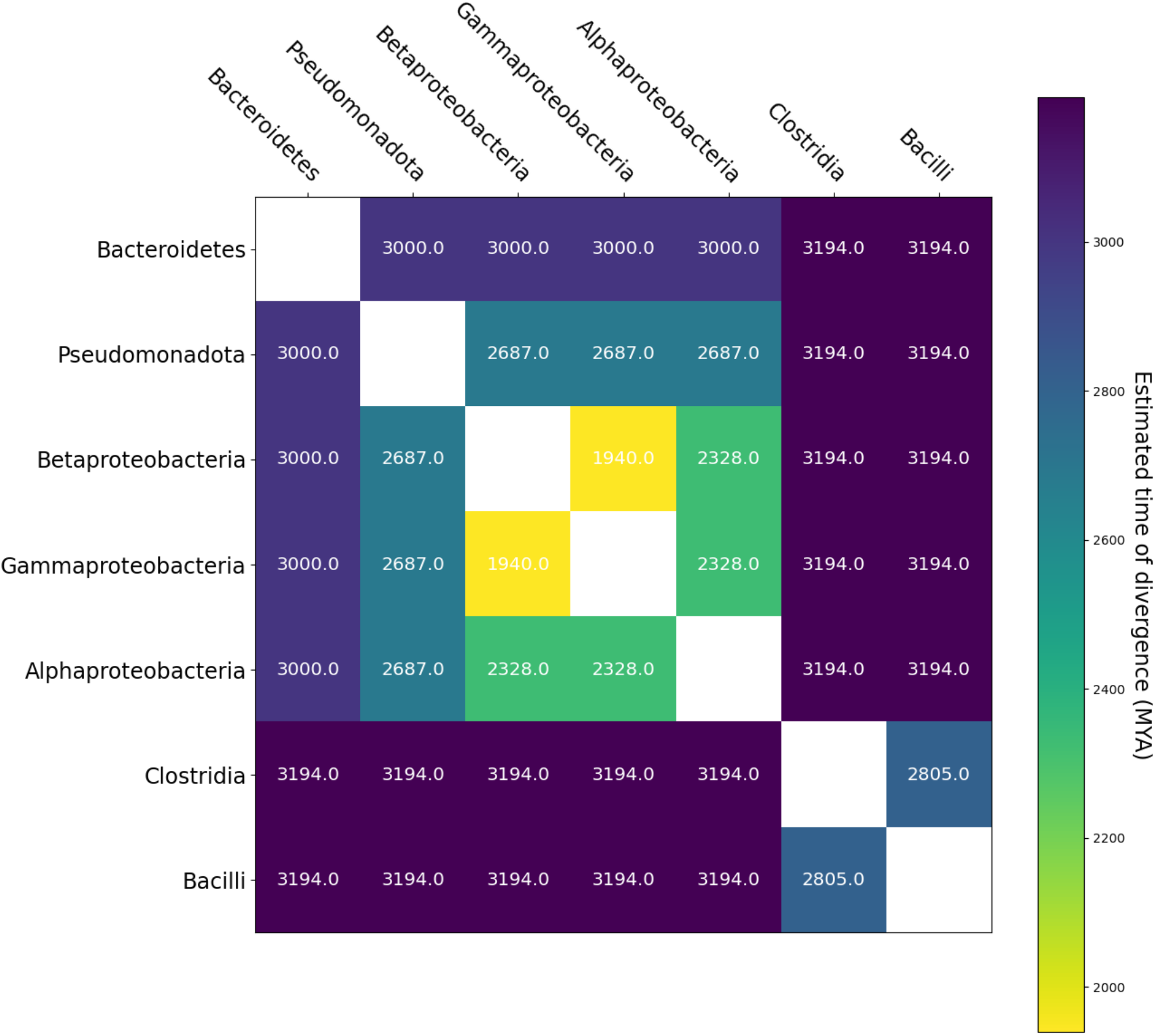
Divergence time between bacterial clades, as estimated by TimeTree (Kumar, et al., 2017). MYA=million years ago.

## Notes

### Competing Interest Statement

The authors have declared no competing interest.

